# Reduced chlorhexidine and daptomycin susceptibility arises in vancomycin-resistant *Enterococcus faecium* after serial chlorhexidine exposure

**DOI:** 10.1101/148809

**Authors:** Pooja Bhardwaj, Amrita Hans, Kinnari Ruikar, Ziqiang Guan, Kelli L. Palmer

## Abstract

Vancomycin-resistant *Enterococcus faecium* (VREfm) are critical public health concerns because they are among the leading causes of hospital-acquired bloodstream infections. Chlorhexidine (CHX) is a bisbiguanide cationic antiseptic that is routinely used for patient bathing and other infection control practices. VREfm are likely frequently exposed to CHX; however, the long-term effects of CHX exposure have not been studied in enterococci. In this study, we serially exposed VREfm to increasing concentrations of CHX for a period of 21 days in two independent experimental evolution trials. Reduced CHX susceptibility emerged (4-fold shift in CHX MIC). Sub-populations with reduced daptomycin (DAP) susceptibility were detected, which were further analyzed by genome sequencing and lipidomic analysis. Across the trials, we identified adaptive changes in genes with predicted or experimentally confirmed roles in chlorhexidine susceptibility (*efrE*), global nutritional stress response (*relA*), nucleotide metabolism (*cmk*), phosphate acquisition (*phoU*), and glycolipid biosynthesis (*bgsB*), among others. Moreover, significant alterations in membrane phospholipids were identified. Our results are clinically significant because they identify a link between serial sub-inhibitory CHX exposure and reduced DAP susceptibility. In addition, the CHX-induced genetic and lipidomic changes described in this study offer new insights into the mechanisms underlying the emergence of antibiotic resistance in VREfm.

## Introduction

*Enterococcus faecium* is a Gram-positive bacterium that naturally colonizes the human gastrointestinal tract and is an opportunistic pathogen associated with bacteremia, urinary tract infections, endocarditis, and wound infections (1-3). Vancomycin-resistant *E. faecium* (VREfm) are of particular concern for infection treatment. VREfm are among the primary etiological agents of central line-associated bloodstream infections (CLABSIs), a type of healthcare-associated infection (HAI) that arises from central venous catheter use and is associated with high mortality in the United States (4, 5). *E. faecium* contamination on indwelling venous catheters, surgical instruments, and hospital surfaces is challenging to eradicate (2, 6-8). In hospital and clinical settings, improper infection control practices, contaminated surfaces, and indiscriminate use of antibiotics contribute to persistence of *E. faecium* (6, 9).

Chlorhexidine (CHX) is a cationic antiseptic and membrane-active antimicrobial (10-12). The primary mechanism of action of CHX is to disrupt the bacterial cell membrane and cause leakage of cytoplasmic contents and precipitation of cytoplasm (13-15). CHX is recommended by the Society for Healthcare Epidemiology of America to reduce CLABSI occurrence in acute care hospitals (16). Specifically, CHX bathing and CHX-impregnated cardiovascular catheters are used for CLABSI control (16-18). Clinical reports have raised concerns about the long-term effects of CHX bathing on hospital-associated pathogens (19-22). The CHX concentrations on patient skin can fall below the MIC for VREfm between bathings (22). Frequent exposure to sub-inhibitory CHX could select for VREfm mutants with reduced susceptibility to CHX and other antimicrobials that also interact with the bacterial cell surface. It was recently reported that colistin resistance emerged in the Gram-negative pathogen *Klebsiella pneumoniae* after exposure to CHX (23).

In a previous study, we used RNA sequencing to study the global transcriptomic responses of a VanA-type VREfm strain to CHX (24). We found that CHX exposure elicited expression of genes associated with antibiotic resistance and extracytoplasmic stress, including genes associated with vancomycin resistance (*vanHAX*) and reduced daptomycin (DAP) susceptibility (*liaXYZ*) (24). In the current study, we test the hypothesis that serial exposure to sub-MIC CHX selects for VREfm mutants with reduced susceptibilities to CHX and other membrane and cell wall-targeting antimicrobials, with particular focus on DAP.

## Materials and Methods

### Bacterial strains and growth conditions

Bacterial strains used in this study are shown in Table 1. *E. faecium* was cultured at 37°C on brain heart infusion (BHI) agar or in BHI broth without agitation unless otherwise stated. Unless otherwise stated, the CHX product used for experiments was Hibiclens (4% wt/vol chlorhexidine gluconate with 4% isopropyl alcohol). Chloramphenicol was used at 15 μg/ml for *E. coli* and *E. faecium*.

**Table 1.**
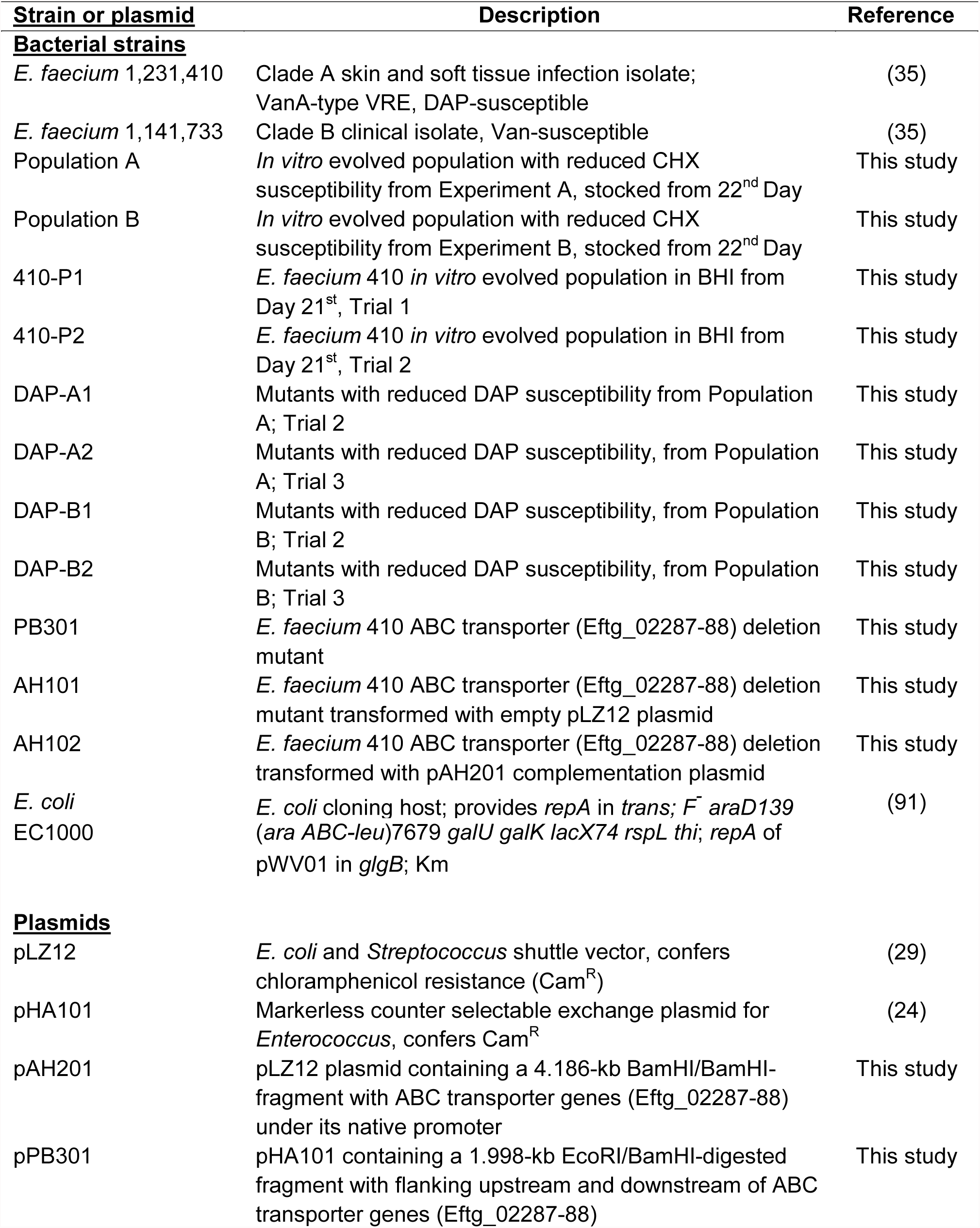
Bacterial strains and plasmids used in the study.

### Routine molecular biology techniques

*E. faecium* genomic DNA (gDNA) was isolated using a previously published protocol (25). DNA fragments were purified using the Purelink PCR purification kit (Thermo Fisher). *Taq* polymerase (New England Biolabs; NEB) was used for routine PCR. Routine DNA sequencing was performed by the Massachusetts General Hospital DNA core facility (Boston, MA). Primers used are shown in Table S1.

### MIC determinations

Vancomycin, ampicillin, and CHX MICs were determined by broth microdilution in BHI as described previously (24). CHX MICs were recorded at 48 hr post-inoculation. DAP MICs were measured using Etest (BioMérieux, Inc.) strips on Mueller-Hinton agar (MHA) plates per the manufacturer’s instructions. MICs were independently assessed at least three times.

### Quantitative RT-PCR (RT-qPCR)

*E. faecium* 1,231,410 (*E. faecium* 410) and 1,141,733 cultures treated for 15 min with 0X (control) and 1X MIC CHX were harvested and mixed with two volumes of RNAProtect Bacteria reagent (Qiagen) according to the manufacturer’s recommendation and a previously published protocol (24). 100 ng RNA was used to synthesize cDNA with Superscript II (Life Technologies) and 5 ng cDNA was used as template in RT-qPCR with primers to amplify internal regions of *liaX* or *clpX*. Threshold cycle (*C_T_*) values were used to calculate the fold change of *liaX* gene expression between 1X MIC CHX-treated cultures and control cultures (n=2 independent trials).

To assess *liaX* expression in the DAP mutants and the *E. faecium* 410 wild-type, overnight cultures inoculated from glycerol stocks were diluted into fresh pre-warmed BHI. The cultures were incubated at 37°C until an optical density at 600 nm (OD_600_) of ~ 0.6 was reached. 15 ml of the cultures were harvested as described above. The experiment was performed independently three times and one-tailed Student’s *t* test was used to assess significance.

### Serial passage experiments

*E. faecium* 410 wild-type was used for *in vitro* serial passage according to a previously published protocol (26).The CHX 1X MIC value for *E. faecium* 410 was 4.9 μg/ml on day 1. For serial passage, overnight culture was adjusted to OD_600_ 0.1 and exposed to CHX below, at, and above the MIC in BHI broth. The cultures were incubated at 37°C, and after 24 hr, the cultures with visible growth in the highest drug concentration were used as inoculum for the next round of passaging on day 2. Again, the inoculum OD_600_ was adjusted to 0.1, and cultures were exposed to CHX as described above. Passages were performed for 21 days. The serial passage experiment was performed independently twice, referred to as Experiments A and B. Cultures from the 22^nd^ day of experiment A (referred to as Population A) and experiment B (referred to as Population B) were analyzed in this study. *E. faecium* 410 cultures were also passaged in BHI without CHX for 21 days in two independent trials. Populations from the 21^st^ day of these trials are referred to as 410-P1 and P2 in this study.

### Agar daptomycin susceptibility assay

Cultures of Population A and Population B were inoculated directly from glycerol stocks into BHI broth and incubated overnight. Cultures were serially diluted in 1X phosphate buffered saline (PBS) and spotted on MHA plates supplemented with calcium (50 μg/ml) with or without 10 μg/ml DAP. Plates were incubated for 24-36 hr at 37°C prior to colony counting. Three (for Population A) or four (for Population B) independent trials were performed. One-tailed Student’s *t* test was used to assess significance. *E. faecium* 410 unpassaged (7 trials) and no-drug passaged populations (3 trials each) were assayed as controls. Colonies arising on DAP plates from two independent trials each for Populations A and B were pooled, inoculated in BHI broth, and cryopreserved.

### Genome sequencing and analysis

Genomic DNA was isolated from overnight broth cultures. DAP MIC was confirmed for these cultures by Etest assay. Library preparation and 2x150 paired end Illumina sequencing was performed by Molecular Research LP (Shallowater, Texas). The *E. faecium* 410 wild-type strain was also sequenced as a control. Sequence reads were first assembled to the previously published *E. faecium* 410 draft genome sequence (NZ_ACBA00000000.1) using default parameters for local alignment in CLC Genomics Workbench (Qiagen). Polymorphisms in the read assemblies were detected using the basic variant mapping tool for sites with ≥10-fold coverage. Variations occurring with ≥50% frequency were compared with the *E. faecium* 410 wild-type read assembly to find mutant-specific SNPs. To detect putative transposon hops, the read mapping parameters were changed to global instead of local alignment, and regions of interest were manually analyzed. ≥98.8% of nucleotide positions in the *E. faecium* 410 reference were covered ≥10-fold and included in variation analyses. Variants unique to the DAP-resistant mutants were confirmed by Sanger sequencing.

### Phosphate assay

A commercially available kit (Sigma MAK030) and a previously published protocol (27) were utilized to measure intracellular inorganic phosphate (Pi) levels. Overnight cultures were diluted to an OD_600_ 0.01 in 50 ml pre-warmed BHI and incubated at 37°C with shaking at 100 rpm. The phosphate levels were measured at four time points, time point 1: OD_600_ 0.4 to 0.5; 2, OD_600_ 0.5 to 0.6; 3, OD_600_ 0.6 to 0.7; 4, OD_600_ 0.7 to 0.8. For each timepoint, 100 μl culture was serially diluted in 0.9% sterile NaCl and spotted on BHI plates for CFU determination. Also, 1 ml of the culture was incubated on ice for 5 min and then pelleted at 13,300 X g for 2 min at 4°C. The pellet was washed twice in 1 ml double-distilled water, resuspended in 0.5 ml double-distilled water, and then disrupted thrice (Fast-Prep-24; MPBio) at 6.5 m/s for 30 s. The homogenized samples were centrifuged at 13,300 X *g* for 15 min at 4°C. Twenty-five μl of the supernatant was diluted with 25 μl double distilled water, and the Pi levels were measured per the manufacturer’s instructions. The phosphate levels were normalized by CFU. One-tailed Student's *t* test was used to assess significance from three independent trials.

### Bacterial growth curves

Overnight cultures from glycerol stocks were diluted to an OD_600_ 0.01 in 50 ml pre-warmed BHI and incubated at 37°C and shaking at 100 rpm. The OD_600_ values were measured for 6 hr. The experiment was performed independently three times and one-tailed Student’s *t* test was used to assess significance.

### Rifampin resistance frequency

Overnight cultures of *E. faecium* 410 wild-type and DAP-resistant mutants were serially diluted in 1X PBS and spot-plated on BHI agar to obtain total CFU counts. Three ml of the cultures were pelleted and spread on BHI agar plates supplemented with 50 μg/ml rifampin to obtain rifampin-resistant CFU counts. Plates were incubated for 24-48 hr. Four independent trials were performed and the significance value was calculated using one-tailed Student’s *t* test.

### Gene deletion and complementation

Loci (EFTG_02287-02288) encoding the predicted ABC transport system EfrEF were deleted in-frame utilizing plasmid pHA101. Briefly, 999 bp flanking upstream and downstream regions of the genes of interest were amplified using primers in Table S1 and ligated with the plasmid pHA101, and propagated in *E. coli* EC1000. The construct, pPB301, was sequence-verified and electroporated into *E. faecium* 410 using a previously published protocol (24). Temperature shift at non-permissible temperature of 42°C and counter-selection with p-chlorophenylalanine was followed according to a previously published protocol (28). Deletion of EFTG_02287-02288 in the mutant strain (*E. faecium* PB301) was confirmed by Sanger sequencing. For complementing this deletion *in trans*, EFTG_02287-02288 with their putative native promoter were amplified and ligated with pLZ12 (29). The complementation construct (pAH201) was sequence-verified and electroporated into the deletion mutant to generate the strain AH102. The empty pLZ12 plasmid was also transformed into the deletion mutant as a control (strain AH101).

### Complementation assay for CHX susceptibility

Overnight cultures of AH101 and AH102 were serially diluted and spot plated on BHI-chloramphenicol (15 μg/ml) plates supplemented with or without 1/8X MIC CHX. The plates were incubated at 37°C for 24-36 hr. The CFU/ml of three independent trials was quantified and one-tailed Student's *t* test was used to assess significance.

### Lipidomic analysis

Overnight cultures from the glycerol stock were inoculated into 10 ml BHI and incubated at 37°C. The 10 ml cultures were added to pre-warmed 250 ml BHI and incubated until an OD_600_ of ~ 0.6 was obtained. 100 μl was removed for DAP Etest testing on MHA plates. Also, 15 ml of the cultures was added to 2 volumes of RNAProtect Bacteria reagent (Qiagen) according to manufacturer’s recommendation and a previously published protocol (24), and samples were used to assess *cls*, *cmk*, *clpX* or *liaX* expression by RT-qPCR as described above. The remaining culture was pelleted at 10,000 rpm at 4°C. Cell pellets were stored at -80°C prior to lipid extraction by the Bligh and Dyer method (30). Lipid analysis by normal phase LC-electrospray ionization (ESI) MS was performed using an Agilent 1200 Quaternary LC system coupled to a high resolution TripleTOF5600 mass spectrometer (Sciex, Framingham, MA), as previously described (31, 32). Normal phase LC was performed on an Agilent 1200 Quaternary LC system using an Ascentis Silica HPLC column, (5 μm; 25 cm x 2.1 mm; Sigma-Aldrich). Mobile phase A and B solvents, flow conditions, and instrumental settings for ESI/MS and MS/MS are as previously published (33). Data analysis was performed using Analyst TF1.5 software (Sciex, Framingham, MA).

### Accession number

Raw Illumina sequencing reads generated in this study is available in the SRA under the accession number SRP108331.

## Results

### *In vitro* evolution of reduced CHX susceptibility in VREfm

Previous RNA sequencing analysis identified up to 118-fold up-regulation of *liaXYZ* in CHX-treated *E. faecium* 1,231,410 (*E. faecium* 410) (24), a VanA-type vancomycin- and ampicillin-resistant blood isolate and member of the hospital-adapted Clade A1 (34, 35). This was of interest because *liaXYZ* is protective against DAP (26, 36-40). We confirmed the CHX-stimulated up-regulation of *liaX* in *E. faecium* 410 and a commensal Clade B strain, *E. faecium* 1,141,733 (35) using RT-qPCR (Fig. S1). We hypothesized that repeat exposure to sub-inhibitory CHX could select for mutants with reduced susceptibility to CHX and concomitant reduced susceptibility to other antimicrobials. To test this, we performed *in vitro* serial passaging of *E. faecium* 410 with CHX for a period of 21 days, starting with a sub-MIC concentration of 2.9 μg/ml (Fig. 1). Similar patterns of MIC shifts were observed for two independent trials over the course of 21 days. The CHX MICs of the evolved populations (referred to as populations A and B) recovered after one drug-free passage was confirmed to be increased (19.6 μg/ml) as compared to the parental strain (Table 2). The CHX MIC was not altered in *E. faecium* 410 passaged for 21 days in medium without CHX. We conclude that reduced CHX susceptibility emerges in VREfm after serial *in vitro* CHX exposure.

**Figure 1.**
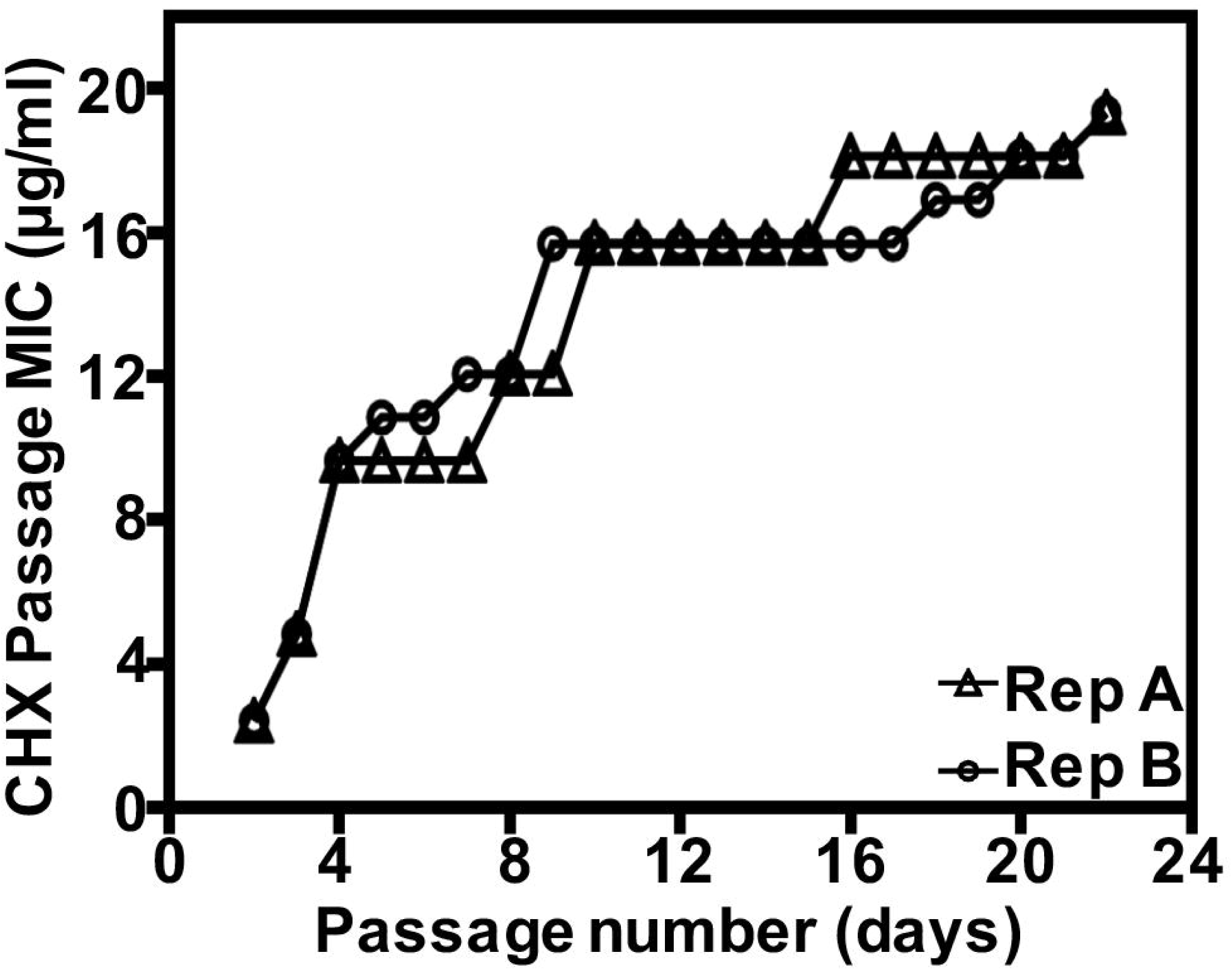
*E. faecium* can adapt to CHX. *In vitro* evolution of reduced CHX susceptibility in *E. faecium* 1,231,410 (*E. faecium* 410) by serial passaging in increasing concentrations of CHX for a period of 21 days. CHX passage MIC (y-axis) for each day of passage (x-axis) is shown for two independent experiments (A and B).

**Table 2.** MIC values for CHX and cell wall-targeting antibiotics.

| Strain or Population | CHX MIC<sup>a</sup> | DAP MIC range, median (*p*-value)<sup>b</sup> | AMP MIC | VAN MIC |
| --- | --- | --- | --- | --- |
| *E. faecium* 410 | 4.9 | 2, 2 (N/A) | 195 | 250 |
| Population A | 19.6 | 2-6, 3 (0.0085) | 390 | 500 |
| Population B | 19.6 | 3-4, 3.5 (0.0006) | 195 | 500 |
| DAP-A1 | 19.6 | 3-6, 4 (0.0043) | 195 | 500 |
| DAP-A2 | 19.6 | 3-6, 4 (0.0015) | 390 | 1000 |
| DAP-B1 | 19.6 | 3-4, 3.5 (0.0006) | 195 | 1000 |
| DAP-B2 | 19.6 | 3-4, 3.5 (0.0006) | 195 | 1000 |

<sup>a</sup>All MICs in table expressed in µg/ml.
<sup>b</sup>DAP MICs were performed independently either 7 (for *E. faecium* 1,231,410 and Population A) or 6 times. The one-tailed unpaired Student's *t* test was used to assess significance of the difference in MIC as compared to *E. faecium* 1,231,410. *E. faecium* 410, *E. faecium* 1,231,410; AMP, ampicillin; Van, vancomycin.

The populations A and B had variable but significantly higher DAP MICs relative to the wild-type strain (Table 2), with the DAP MIC in some experimental trials meeting the ≥4 μg/ml breakpoint for DAP resistance (41). Vancomycin MIC was 2-fold higher for both populations relative to wild-type, and ampicillin MIC was 2-fold higher for population A.

### Reduced DAP susceptibility is present in the CHX-passaged populations

We sought to further quantify and investigate the basis for elevated DAP MIC in the CHX-passaged populations. The CHX-passaged populations (A and B) were cultured on agar with and without 10 μg/ml DAP to quantify CFU (Fig. 2). *E. faecium* 410 wild-type stock cultures (410-1 and -2) and *E. faecium* 410 passaged for 21 days in the absence of CHX (410-P1 and -P2) were used as controls. Sub-populations with reduced DAP susceptibility were detected in the CHX-passaged populations (Fig. 2).

**Figure 2.**
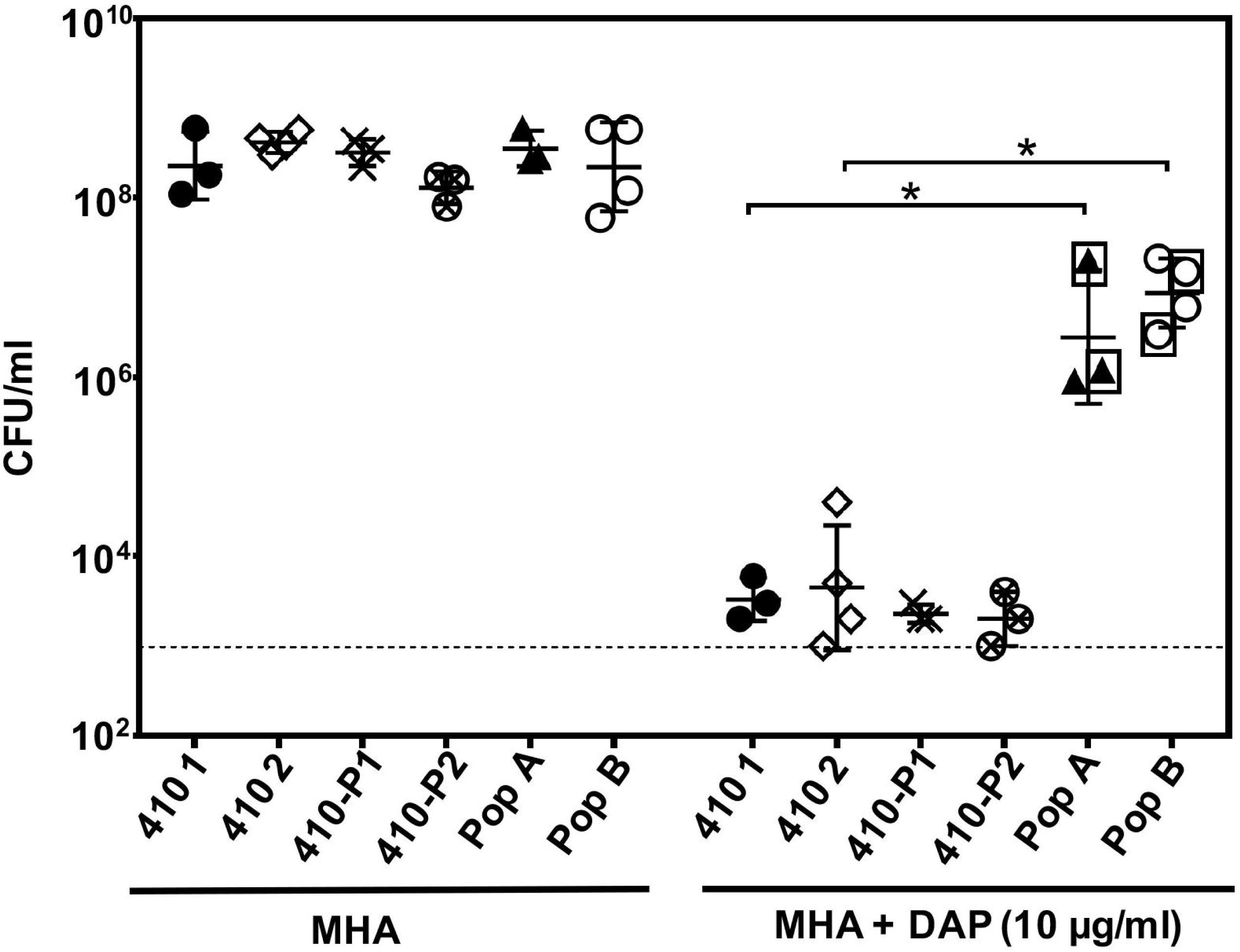
Reduced DAP susceptibility in CHX-passaged *E. faecium* populations A and B. The geometric mean and geometric standard deviation of CFU/ml count for n=3 or 4 independent trials is shown. Strains and populations are described in the text. The dashed line represents limit of detection (10^3^ CFU/ml). The boxed populations were sent for whole genome sequencing. *, *P* < 0.05; one-tailed Student’s *t* test.

Next, isolated colonies arising on DAP plates from each of two independent DAP plating trials were pooled and stocked for further analysis. These mutants will be referred to as DAP-A1 and DAP-A2 (from population A; 14 and 11 colonies were pooled, respectively) and DAP-B1 and DAP-B2 (from population B; 18 and 5 colonies were pooled, respectively) hereafter. The DAP MICs of these mutants after two DAP-free passages were found to be variable but significantly higher than the *E. faecium* 410 wild-type (Table 2). The DAP mutants, with the exception of DAP-B2, have significantly longer generation times than *E. faecium* 410 wild-type in BHI broth (Table S2). No significant differences in spontaneous rifampin resistance frequencies were observed between the DAP mutants and *E. faecium* 410 wild-type (Table S2), indicating that these strains are not hyper-mutators. RT-qPCR analysis identified a small but statistically significant increase in *liaX* expression in the DAP-A1 and A2 mutants relative to wild-type. The increase was insignificant for DAP-B1 and B2 (Fig. S2).

### DAP strains have mutations in genes previously associated with reduced antimicrobial susceptibilities

Genome sequencing identified mutations occurring in the DAP strains relative to *E. faecium* 410 wild-type (Table 3). All four share a common mutation in *efrE* (EFTG_02287), which encodes one subunit of the heterodimeric ABC transporter EfrEF (42). *efrE* is up-regulated 22-fold by *E. faecium* 410 in response to CHX (24), and deletion of *efrE* from *E. faecalis* OG1RF confers increased susceptibility to CHX (43, 44). No mutations other than in *efrE* were common to all strains.

**Table 3.** List of mutations identified by whole-genome sequencing in the DAP-resistant mutants.

| Mutant | Description of gene | Nucleotide variation (frequency)<sup>a</sup> | Amino acid<br>change | *E. faecalis* V583<br>ortholog |
| --- | --- | --- | --- | --- |
| **DAP-A1** | Eftg_02287 ABC transporter<br>Eftg_01724 RelA/SpoT (*relA*)<br>Eftg_01135 Alpha/beta hydrolase<br>Eftg_00577 Cytidylate kinase (*cmk*)<br>promoter | C67008T (98.47%)<br>CTAGGATTTAC30479Del (98.19%)<br>G137748A (55.64%)<br>A618699IS 1251<sup>b</sup> | Ala290Val<br>Leu505fs<br>Trp150*<br>IS 1251 insertion | EF2226<br>EF1974<br>EF1505<br>EF1547 |
| **DAP-A2** | Eftg_02287 ABC transporter<br>Eftg_01724 RelA/SpoT (*relA*)<br>Eftg_01135 Alpha/beta hydrolase<br>Eftg_00577 Cytidylate kinase (*cmk*)<br>promoter | C67008T (99.77%)<br>CA29139AG (95.74%)<br>137847G insertion (88.17%)<br>A618699IS 1251<sup>b</sup> | Ala290Val<br>Ala58Glu<br>Ile184fs<br>IS 1251 insertion | EF2226<br>EF1974<br>EF1505<br>EF1547 |
| **DAP-B1** | Eftg_02287 ABC transporter<br>Eftg_01175 PhoU regulator<br>Eftg_02534 Transposase | C67008T (98.962%)<br>G173845A (99.61%)<br>C71844A (99.47%) | Ala290Val<br>Glu61Lys<br>Lys206Asn | EF2226<br>EF1754 |
| **DAP-B2** | Eftg_02287 ABC transporter<br>Eftg_02162 Glycosyltransferase (*bgsB*) | C67008T (99.36%)<br>C50223T (67.09%) | Ala290Val<br>Val57Met | EF2226<br>EF2890 |

<sup>a</sup>Frequency of mutation in the read assembly as determined by variant detection in CLC Genomics Workbench.
<sup>b</sup>Precise location of insertion was mapped by Sanger sequencing.

DAP-A1 and DAP-A2 share IS*1251* insertions in the promoter region of *cmk*, which encodes cytidylate kinase (Table 3). RT-qPCR confirmed the down-regulation of *cmk* in DAP-A2 in the two trials but not in DAP-A1 (Fig. S3). Population heterogeneity in regards to IS*1251* insertion at the *cmk* promoter was observed in DAP-A1 by gel electrophoresis analysis of *cmk* promoter amplicons (data not shown).

DAP-A1 and DAP-A2 each have different mutations in *relA* and in a gene encoding a predicted alpha/beta hydrolase (Table 3). Production of (p)ppGpp, a bacterial alarmone, is controlled by RelA. Changes in ppGpp concentration modulate the stringent stress response and impact antibiotic tolerance and virulence in *Enterococcus* (45, 46). Mutations in *relA* have previously been associated with *in vitro* (47) and *in vivo* (48) emergence of DAP resistance in *Bacillus subtilis* and *E. faecium*, respectively. In DAP-A1, deletion of 11 bases (CTAGGATTTAC) from *relA* results in synthesis of a truncated, 505 amino acid protein that lacks the C-terminal ACT regulatory domain. The function of ACT domain as suggested is that it regulates the catalytic activities of the amino terminal domain to ensure that synthesis and degradation of ppGpp are not co-stimulated (49, 50). In DAP-A2, the A58E substitution is predicted by EMBOSS (51) to convert a beta-strand fold into an alpha helix in the HD4 metal-dependent phosphohydrolase domain of RelA. DAP-A1 and DAP-A2 also possess nonsense and frameshift mutations, respectively, in a gene encoding a predicted alpha/beta hydrolase. To our knowledge, this gene has not previously been linked with antimicrobial susceptibility.

Different mutations were identified in DAP-B1 and DAP-B2. In DAP-B1, a E61K substitution occurs in PhoU. PhoU regulates phosphate intake by high-affinity inorganic phosphate-specific transporters (*pst* transporters encoded by EFTG_01170-74 in *E. faecium*) in *E. coli* (52-56). The *pst* transport system is up-regulated in *Streptococcus pneumoniae* in response to penicillin (57), and mutations in the *pst* system led to decreased accumulation of reactive oxygen species after exposure to penicillin (58). Moreover, an adaptive point mutation in a putative phosphate transporter gene (*pitA*) in *S. aureus* conferred growth phase-dependent tolerance to DAP (27). However, the exact molecular mechanism(s) of how inorganic phosphate levels alter susceptibility to these antimicrobials is unclear. We confirmed that inorganic phosphate concentrations were significantly lower in the DAP-B1 mutant relative to the wild-type for two of four time-points assayed (Fig. S4).

In DAP-B2, a V57M substitution occurs in BgsB (59). BgsB, a glycosyltransferase (GT), catalyzes the transfer of glucose from UDP-glucose to 1,2-diacylglycerol (DAG) forming 3-D-glucosyl-1,2-diacylglycerol (MGlcDAG). Further, MGlcDAG is converted to diglucosyl DAG (DGlcDAG) by BgsA (60). MGlcDAG and DGlcDAG are important glycolipids for cell membrane fluidity and lipoteichoic acid (LTA) synthesis. Deletion of *bgsB* in *E. faecalis* 12030 led to a complete loss of cell membrane glycolipids and affected the chain length of glycerol phosphate polymer of LTA. It also led to 2-fold increased sensitivity to antimicrobial peptides (colistin and polymixin B), and reduced virulence in a rat model of endocarditis (59, 61). Deletion of other cell wall glycosyltransferases (*epaOX* and *epaI*) in *E. faecalis* enhances susceptibility to DAP (62) through perturbations in the cell envelope. A role for GTs in the production of immature polysaccharides, which prevented DAP binding, was suggested.

### *efrE* impacts CHX susceptibility

Because *efrE* mutations were shared by all DAP strains, we deleted the EfrEF transport system from *E. faecium* 410 to assess its role in CHX and DAP susceptibility. Deletion of *efrEF* resulted in a 4-fold decrease in CHX MIC as compared to *E. faecium* 410 (n=3 independent trials). The median DAP MIC of the Δ*efrEF* mutant was determined to be 2 μg/ml (range, 2-3 μg/ml; n=5 independent trials; *p*-value = 0.09 compared to *E. faecium* 410 wild-type using one-tailed Student’s *t* test). Complementation *in trans* with *efrEF* on a multi-copy vector (AH102) resulted in a 2- to 4-fold increase in CHX MIC as compared to an empty vector control strain (AH101) using broth microdilution assays (n=2 independent trials). However, we observed a growth inhibition effect when chloramphenicol was added to the broth microdilution assay for vector selection. Hence, we utilized a spot-plating assay to quantify CFU differences between strains AH101 and AH102 in the presence of CHX and chloramphenicol (Fig. S5). Complementation of ABC transporter genes *in trans* resulted in significant increase in CFU count in the presence of CHX compared to control cultures.

To identify a putative function for EfrEF, we performed comparative lipidomic analysis of *E. faecium* 410 and the *efrEF* deletion mutant. The most striking differences observed were for two species whose positive ion [M+H]^+^ signals are detected at m/z 288 and m/z 316 by ESI/MS. These two species are present in *E. faecium* 410 wild-type, but absent in the *efrEF* deletion mutant (Fig. S6). High resolution mass measurement and tandem MS analysis identified these species as ethoxylated fatty amines previously identified as components of the anti-static additive Atmer-163 (63) commonly used in consumer products such as plastics. The *m/z* 288 and 316 ions correspond to the [M+H]^+^ ions of Atmer-163 containing C_13_ and C_15_ fatty alkyl chains, respectively. The absence of Atmer-163 (C_13_) and Atmer-16 (C_15_) in the *efrEF* mutant suggests that this ABC transporter is involved in transporting the Atmer-163 species, with lipid-like structures, from media into the cells.

### Distinct alterations of membrane phospholipid compositions in DAP mutants

We next compared lipid profiles between the DAP mutants and *E. faecium* 410 wild-type. The lipid extracts were subjected to normal phase LC-ESI/MS using a silica column for lipid separation. As shown by the total negative ion chromatogram data in Fig. 3 and Table S3, cardiolipin (bisphophatidylglycerol; CL), an anionic phospholipid, was dramatically reduced in DAP-A mutants (10-fold in DAP-A1 and 12-fold in DAP-A2) as compared to *E. faecium* 410. As expected, an increased accumulation of the precursor phosphatidic acid (PA) (4-fold in DAP-A1 and 5-fold in DAP-A2) was also detected. The expression levels of two predicted cardiolipin synthase (*cls*) genes (EFTG_00614 and EFTG_01168), which mediate reversible transphosphatidylation of PG molecules to synthesize cardiolipin, were not significantly altered compared to *E. faecium* 410 (data not shown). In contrast to DAP-A mutants, CL and PA contents were not significantly altered in DAP-B mutants (Table S3), suggesting that DAP-A and DAP-B mutants do not share identical molecular mechanisms for reduced DAP susceptibility despite their similar DAP phenotypes.

**Figure 3.**
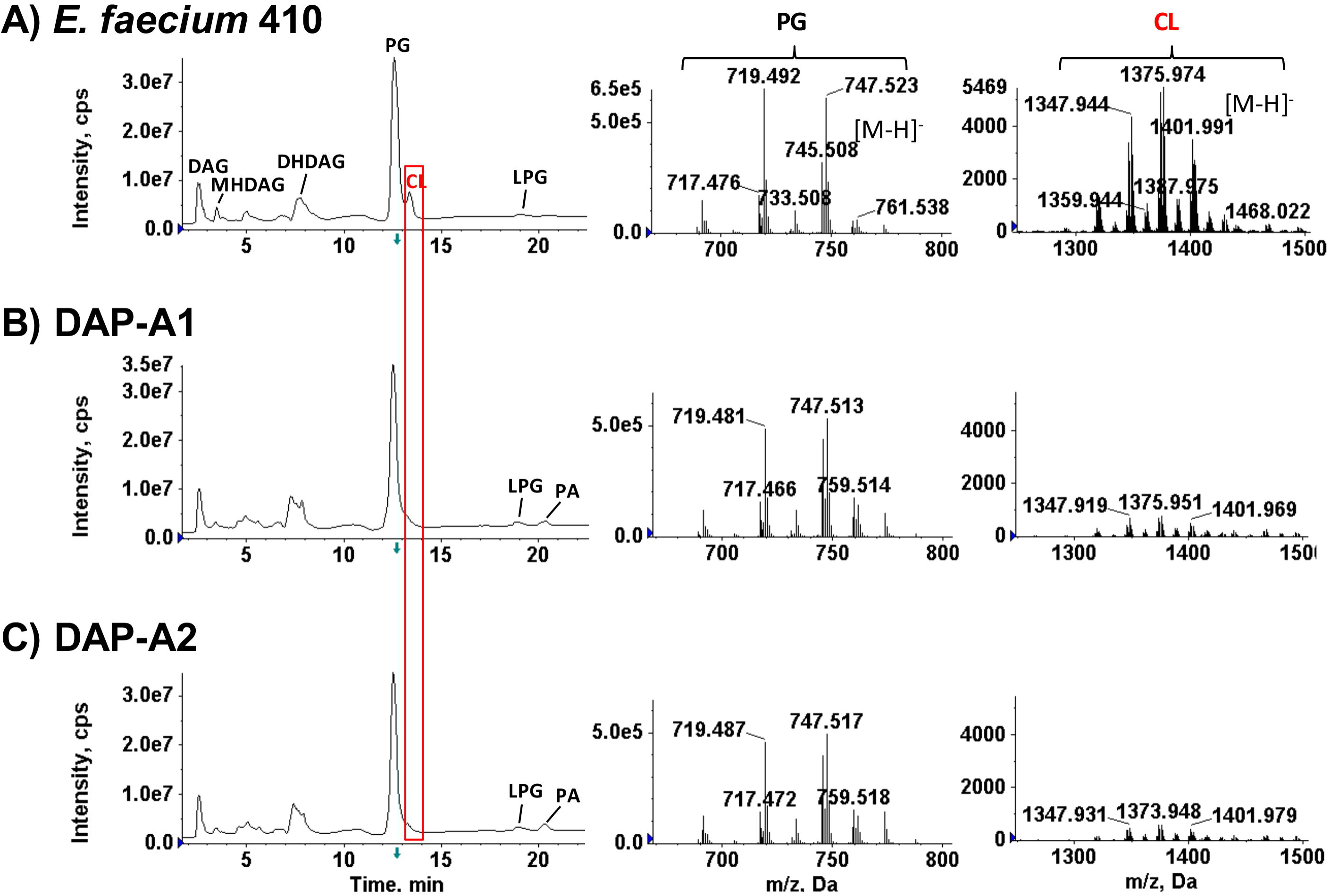
Cardiolipin levels are decreased in DAP-A1 and DAP-A2 mutants. Lipidomic analysis of *E. faecium* 410 and DAP strains DAP-A1 and -A2 was performed by normal-phase LC-ESI/MS in the negative ion mode. The major lipids detected are phosphaditylglycerol (PG), cardiolipin (CL), diacylglycerol (DAG), monohexosyldiacylglycerol (MHDAG), dihexosyldiacylglycerol (DHDAG), phosphatidic acid (PA), and lysylphosphaditylglycerol (LPG). While the PG levels are similar in all three strains, the CL levels are significantly decreased and PA are increased in DAP-A1 and -A2 mutants compared to *E. faecium* 410. Shown are the total ion chromatograms (TIC) and the selected mass spectra of [M-H] ion species of PG and CL.

Significant changes were observed in several other lipid species in DAP mutants. As shown in Table S3, in DAP-B1, levels of GPDD (glycerolphosphate DGlcDAG), a LTA precursor, were about 4-fold and lysylphosphaditylglycerol (LPG) were 4-fold higher than *E. faecium* 410 wild-type. In DAP-B2, amounts of monoglucosyl DAG (MGlcDAG or MHDAG) were 2-fold higher, potentially as a result of the *bgsB* mutation. The levels of GPDD were at least 2-fold higher for all DAP-A and -B strains.

## Discussion

The goal of this study was to test the hypothesis that serial exposure to sub-MIC CHX selects for VREfm mutants with reduced susceptibilities to CHX, with concomitant effects on susceptibility to other membrane and cell wall-targeting antimicrobials. Our serial passage experiments demonstrate that reduced CHX susceptibility can emerge in VREfm after repeat sub-inhibitory exposure. Moreover, our DAP plating experiments demonstrate that reduced DAP susceptibility concomitantly emerges in a sub-population of CHX-passaged cells. Using genomics and lipidomics, we identified genetic and physiological changes occurring in these sub-populations with reduced DAP susceptibility.

CHX, a cationic antiseptic, interacts with the bacterial cell membrane (12). Various mechanisms of reduced CHX susceptibility in Gram-negative and -positive bacteria have been reported. The two main mechanisms include CHX efflux (23, 42-44, 64-67) and changes in outer membrane content (68, 69). The recently identified two-component system ChtRS contributes to CHX tolerance in *E. faecium*, presumably via regulation of expression of genes in its regulon, which is currently undefined (70). In this study, we have confirmed a role for the heterodimeric ABC transporter in CHX susceptibility in VREfm. Deletion of *efrEF* increases susceptibility of *E.faecium* 410 to CHX, and an amino acid substitution in EfrE is associated with decreased susceptibility to CHX. The substrate of EfrEF was not assessed in prior studies. By lipidomic analysis, we found that presence of two ethoxylated fatty amine fatty alkyl diethanolamine compounds was abolished in the *efrEF* deletion mutant relative to wild-type. It remains to be determined if these compounds are protective against CHX. Nonetheless, their lipid-like structures may provide useful clues in identifying the molecular species that EfrEF transports, or in the functional studies of EfrEF.

Daptomycin (DAP) is a cyclic lipopeptide antibiotic used to treat infections caused by multidrug-resistant Gram-positive pathogens including VREfm (71-73). DAP resistance arises by mutation, leading to treatment failure (74). DAP is a negatively charged molecule that requires calcium ions for activity. Interaction of the cationic DAP-calcium complex with the membrane induces daptomycin oligomerization, membrane phospholipid remodeling, and other physiological alterations, ultimately leading to cell death (47, 75-79). Broadly speaking, alterations in cell surface composition and in cellular stress responses are associated with reduced DAP susceptibility in Gram-positive bacteria (80, 81).

In this study, we identified adaptive changes in genes with predicted or experimentally confirmed roles in chlorhexidine susceptibility (*efrE*), global nutritional stress response (*relA*), nucleotide metabolism (*cmk*), phosphate acquisition (*phoU*), and glycolipid biosynthesis (*bgsB*) occurring in the CHX-passaged mutants with reduced DAP susceptibilities. That the strains arising on DAP plates are not hyper-mutators indicates that these mutations arose as a result of CHX selection. It remains to be determined at what point in the CHX serial passage experiments reduced DAP susceptibility emerged, as only the beginning and end-points of the evolution experiments were assessed in this study. Deep sequencing of populations at beginning, mid- and end-points of the CHX passage experiments could be used in future studies to further examine the diversity and frequency of genetic variations arising as a result of serial sub-inhibitory CHX exposure. Moreover, it remains to be determined whether susceptibility to host-associated cationic antimicrobial peptides is also altered as a result of serial CHX exposure.

Lipidomic analysis of the DAP mutants identified major alterations in membrane lipid compositions, underscoring a link between membrane lipid compositions and DAP susceptibility. CL, an anionic phospholipid, is present at the septal and polar regions in the bacterial cell membrane and is important for the regulation of processes like cell division, membrane transport, and localization of proteins to specific sites (82, 83). Since DAP preferentially binds to charged regions of the membrane, we hypothesize that reduced amounts of CL in DAP mutants from Population A can affect the binding of DAP antibiotic to the membrane regions. Tran et al showed that remodeling of CL can divert DAP away from division septum in *E. faecalis* (84). In Population B, we detected elevated levels of MHDAG and LTA precursor, implying that LTA synthesis is impaired in these DAP strains. Zorko et al showed direct binding and inhibition of LTA synthesis by CHX using fluorescence displacement and isothermal titration (85). Another study showed inactivation of LTA by CHX in *E. faecalis* (86). DAP mechanism of action by inhibition of LTA synthesis has also been suggested (87, 88) but recent reports have not supported this mechanism (89, 90). In future studies, we will assess the specific contributions of individual mutations identified in the DAP mutants to the lipid content alterations observed.

Our work has clinical implications. If sub-inhibitory CHX exposure selects for VREfm mutants with enhanced abilities to tolerate or resist DAP, these mutants could contribute to treatment failures with DAP. Frequent improper use of CHX (i.e., presence of sub-inhibitory concentrations on patient skin) may favor the emergence and persistence of these VREfm mutants in healthcare settings. Surveillance of VREfm from hospital wards utilizing CHX bathing would be useful to monitor the long-term impact of CHX bathing on these organisms. Routine sub-inhibitory CHX exposure may be a contributing factor to the clinical emergence of DAP resistance in VREfm.

## Acknowledgements

We would like to thank Dr. Ronda Akins for providing daptomycin. This work was supported by start up funds from the University of Texas at Dallas to K.L.P. The MS facility in the Department of Biochemistry at Duke University Medical Center and Z.G. were partially supported by grants GM-069338 and EY023666 from the National Institutes of Health.

## Supplemental Figures and Tables

**Figure S1. CHX induces *liaX* gene expression in *E. faecium***. RT-qPCR was used to quantify the expression of *liaX* upon exposure to CHX for 15 min as compared to control untreated condition. Expression of *liaX* was internally normalized to *clpX* and expression in control cultures was set to 1 (not shown). The fold change in *liaX* expression in cultures treated with 1X MIC CHX relative to the control was quantified for two independent (Trial 1 and 2) experiments. Efm 410, *E. faecium* 1,231,410; Efm 733, *E. faecium* 1,141,733.

**Figure S2. *liaX* gene expression in DAP strains vs. parental strain**. RT-qPCR was used to quantify the expression of *liaX* in the DAP strains at exponential phase (OD_600_~0.6) vs. *E. faecium* 410 wild-type in three independent trials. Expression of *liaX* was internally normalized to *clpX* and *liaX* expression in control cultures was set to 1. The standard deviation was calculated from n=3 independent experiments and one-tailed Student’s *t-*test was used to calculate significance value. *, *P* < 0.05. 410 wild-type, *E. faecium* 1,231,410.

**Figure S3. RT-qPCR for quantifying the expression levels of cytidylate kinase (*cmk*).** RTqPCR was used to quantify the expression of *cmk* in DAP mutant DAP-A1 and DAP-A2 vs. *E*. *faecium* 410 during exponential growth (OD_600_ ~ 0.6). Expression of *cmk* was internally normalized to *clpX*. Expression of *E. faecium* 410 *cmk* was set to 1 (not shown). The fold change in *cmk* expression was quantified for two independent experiments.

**Figure S4. Quantification of intracellular organic phosphate (Pi) levels in *E. faecium* 410 wild-type and DAP-B1 mutant.** Intracellular Pi levels were measured for wild-type and DAP strain DAP-B1 at different growth time points (OD_600_ 0.4-0.8) as described in materials and methods. The levels (pmoles) were normalized using CFU count. Standard deviation was calculated from n=3 independent experiments and significance value was calculated using one-tailed Student’s *t* test. Time points: 1, OD_600_ 0.4-0.5; 2, OD_600_ 0.5-0.6; 3, OD_600_ 0.6-0.7 and 4, OD_600_ 0.7-0.8. *, *P* < 0.05. 410 wt, *E. faecium* 1,231,410.

**Figure S5. Complementation of** Δ***efrEF* deletion mutant.** The average CFU count of the Δ*efrEF* deletion mutant transformed with empty pLZ12 vector and complementation vector pAH201 on BHI and chloramphenicol plate supplemented with or without 1/8X MIC levels of CHX from n=3 independent biological trials is shown. The CFU count was comparable for two strains on BHI plate supplemented with chloramphenicol. A significantly higher CFU count of *efrEF* deletion mutant transformed with complementation vector was observed. *, *P* < 0.05 value was calculated using one-tailed Student’s *t* test. Cam, chloramphenicol.

**Figure S6. Lipid-like compounds (Atmer-163) are detected in *E. faecium* 410 but not in** Δ***efrEF* mutant. A)** Positive ion ESI mass spectra showing the detection of Atmer-163 (C_13_) ([M+H]^+^ at *m/z* 288) and Atmer-163 (C_15_) ([M+H]^+^ ion at *m/z* 316) in *E. faecium* 410, and their absence in the Δ*efrEF* mutant. **B**) Chemical structures and molecular formulae of Atmer-163 (C_13_) and Atmer-163 (C_15_). **C**) MS/MS spectrum of Atmer-163 (C_15_) [M+H]^+^ ion at *m/z* 316. The fragment ion structures are depicted.

**Table S1.** List of primers used in the study.

**Table S2.** Doubling time and Rifampin mutation frequency for the DAP strains vs. *E. faecium* 410 wild-type.

**Table S3.** The relative levels of major lipids identified in *E. faecium* 410 wild-type and DAP strains.

